# Nova-ST: Nano-Patterned Ultra-Dense platform for spatial transcriptomics

**DOI:** 10.1101/2024.02.22.581576

**Authors:** Suresh Poovathingal, Kristofer Davie, Roel Vandepoel, Nicholas Poulvellarie, Annelien Verfaillie, Nikky Corthout, Stein Aerts

**Author notes:** These authors contributed equally (Suresh Poovathingal, Kristofer Davie).

## Abstract

Spatial transcriptomics workflows using barcoded capture arrays are commonly used for resolving gene expression in tissues. However, existing techniques are either limited by capture array density or are cost prohibitive for large scale atlasing. We present Nova-ST, a dense nano-patterned spatial transcriptomics technique derived from randomly barcoded Illumina sequencing flow cells. Nova-ST enables customized, low cost, flexible, and high-resolution spatial profiling of large tissue sections. Benchmarking on mouse brain sections demonstrates significantly higher sensitivity compared to existing methods, at reduced cost.

**Motivation:** Spatial transcriptomics analysis is becoming exceedingly important in biomedical and clinical research. Several platforms for spatial transcriptomics are currently available, but most of these technologies are commercialized making them inflexible and cost prohibitive. The motivation for this work was to establish an open source, flexible and sensitive sequencing-based spatial transcriptomics platform that can provide a considerable cost advantage for performing large scale spatial profiling. We provide thorough and in-depth guidance and resources both for the experimental and computational components of the workflow, to facilitate easy implementation of Nova-ST.

## Introduction

Characterizing and modeling complex tissues requires an understanding of the spatial cellular organization and, of the interactions between cells in the context of normal and pathological states^1,2^. Multiplexed in situ imaging-based assays measure RNA expression at sub-cellular resolution, but their implementation is non-trivial and requires pre-selected gene panels for measurements^3^. Compared to in situ methods, spatial barcoding methods based on oligonucleotide arrays are straightforward, are often easier to implement and, enable unbiased whole transcriptome analysis, Spatial Transcriptomics (currently, 10X Genomics Visium & recently released Visium HD)^4,5^ is one such method that is widely used for spatial RNA sequencing. Other sequencing-based spatial assays use randomly barcoded bead layers for RNA capture (commercialized by Curio Biosciences)^6^ or microfluidic technology to perform deterministic spatial barcoding^7^. Most of these methods are however limited by low spatial resolution, where the active capture area cannot achieve single cell resolution.

Recently, several methods based on nano-patterned arrays have emerged that can potentially provide whole transcriptome capture at subcellular resolution^8–10^. These also allow fine tuning of the size of the capture array to achieve near single cell resolution. Stereo-seq from BGI STomics uses randomly barcoded DNA nanoballs captured on a nanopatterned array^8^. Alternatively, Seq-Scope, repurposes the Illumina MiSeq flow cell to perform spatial RNA sequencing^10^. In the latter method, spatial barcoding is achieved using local bridge amplification of DNA libraries containing random spatial barcodes^10^. In Seq-Scope, the functionalized area of the flow cell where the tissue is overlayed for spatial sequencing is quite limited (< 2mm of imaging area). In addition, MiSeq flow cells have surface functionalization in a contiguous circular pattern (blue numbered circles in Figure 1A), further decreasing the effective spatial footprint, and limiting the size of tissues able to be profiled. With a single MiSeq flow cell, it is only possible to perform spatial sequencing of a total of ∼10-14 mm^2^, meaning that for most practical purposes related to large scale, high throughput spatial sequencing of tissues required for cell-atlasing efforts, or for large sectioned clinical samples, Seq-Scope offers very limited possibilities.

**Figure 1:**
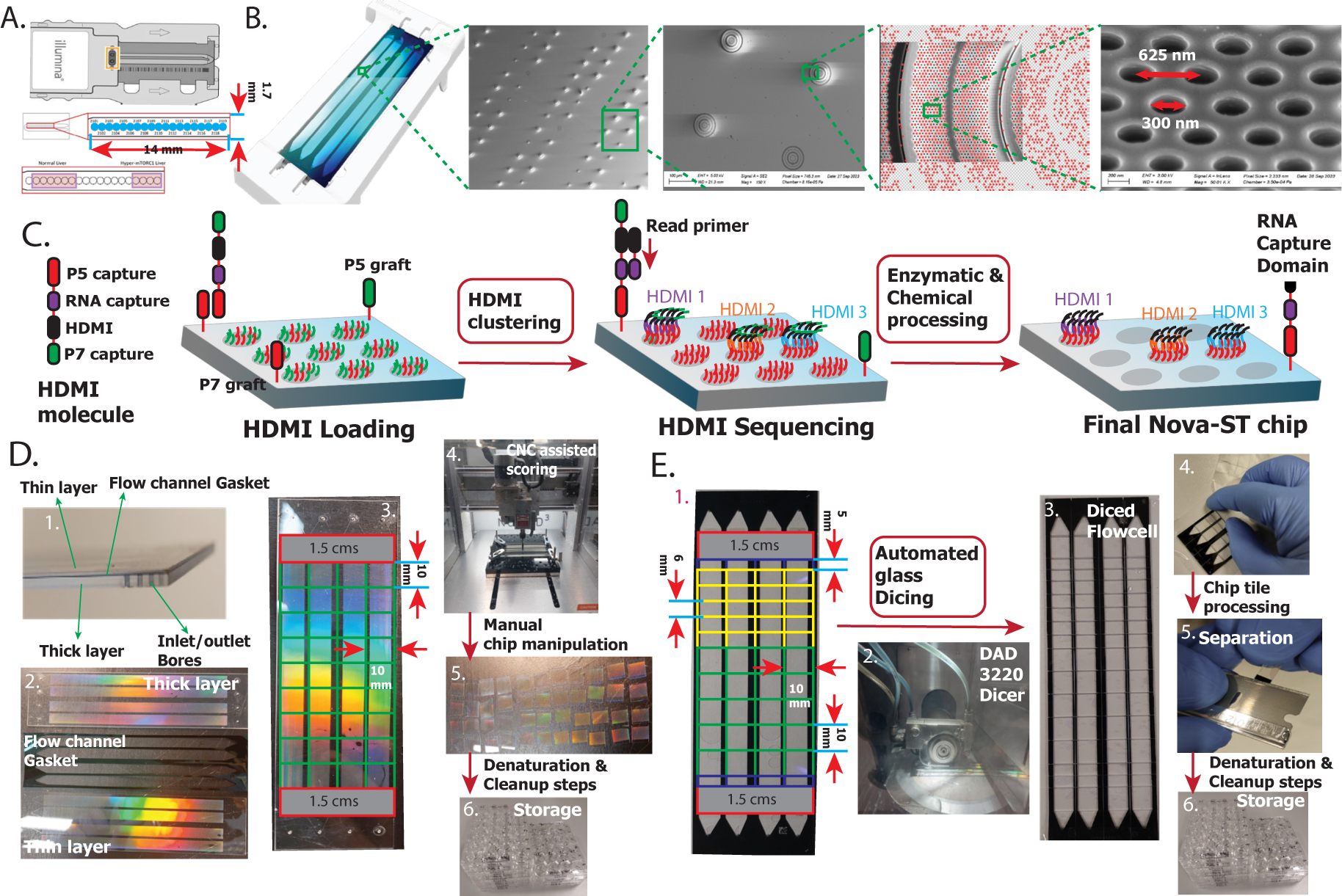
Workflow for Nova-ST chip preparation. A. Details of MiSeq flow cell used in the Seq-Scope platform^10^. Expanded view of the MiSeq’s flow channel indicating the functional area for HDMI capture (the contiguous circular region shaded blue is where the flow cell is functionalized for HDMI barcoding). B. Details of the functionalized surface of NovaSeq 6000 S4 sequencing flow cell. Electron microscopy details of the nanowell pattern on the functional surface of the sequencing flow cell. C. Zoomed in illustration of the nano wells showing the HDMI sequencing on the NovaSeq flow cell and the finalized Nova-ST chip after all the enzymatic and chemical processing resulting in single stranded DNA with HDMI sequences and RNA capture domains. D. Post processing details after the HDMI sequencing for manual cutting: 1. Details of the NovaSeq glass chip showing the thin and thick glass layers sandwiched by a flow gasket. 2. Separated layers of NovaSeq 6000 flow cell exposing the functional surface of the sequencing flow cell. 3. 1 cm x 1 cm grid along which the flow cells are cut to prepare the Nova-ST chips. 1.5 cm thick regions near the entrance and the exit of the flow cell are not sequenced by the NovaSeq 6000 and this region is either discarded or kept for optimizations (grey region). 4. Scribing of the Nova-ST glass layers using the NOMAD 3 CNC milling machine. 5. Final cut 1cm x 1 cm Nova-ST chips. 6.Storage of Nova-ST chips in 24 well plates. E. Post processing details after the HDMI sequencing for automated cutting: 1. A grid pattern of various sizes, along which the flow cells are cut to prepare the Nova-ST chips. 2. Cutting the flow cell into chips using the DAD 3220 Dicer. 3. The diced flow cell, still attached to a dicing tape that keeps the chips together. 4. Removal of the chips from the backing. 5. Separating the thin and thick layers using a sharp razor blade 5. Storage of Nova-ST chips in 24 well plates.

In this work, we developed Nova-ST, a cost effective and easy to use spatial transcriptomics platform to perform spatial sequencing on large tissue sections using high density, patterned flow cells from Illumina. After assessing different flow cells, we chose the NovaSeq 6000 S4 flow cells as they provide the largest area for spatial sequencing amounting roughly to 6000 mm^2^, while also providing a cost-effective sequencing kit with minimal waste for barcoded array generation. Nova-ST consists of both an experimental and computational workflow. Guidance & resources needed for implementing and performing the Nova-ST experimental workflow are available at protocols.io and https://nova-st.aertslab.org/, while the computational pipeline is available at https://github.com/aertslab/Nova-ST.

## Results

### Generation of spatial transcriptomics arrays for Nova-ST

NovaSeq flow cells are patterned with oligonucleotide-functionalized nano-wells arranged in a hexagonal pattern (Figure 1B). Compared to MiSeq flow cells, which are not patterned, the nano-functionalized surfaces of NovaSeq flow cells have several benefits including higher spatial density and lower cluster crosstalk. We imaged the functionalized surface of a NovaSeq 6000 S4 flow cell with electron microscopy (EM), which revealed nano-wells of ∼300 nm diameter and a well-to-well pitch of ∼625 nm (Figure 1B), providing a resolution of ∼350 nano-wells spots for RNA capture per 100 µm^2^ area. Similar to Seq-Scope^10^, our Nova-ST workflow starts with sequencing of High Density Molecular Indexing (HDMI) oligonucleotides. HDMI molecules contain Illumina sequencing adapters, a 32-base long randomer spatial identifier sequence and a DraI restriction enzyme cleavable RNA capture site (Figure 1C, Table S1). The spatial coordinates on the flow cells were created during the first sequencing pass where clonal clusters of HDMI molecules are generated by local cluster amplification of a single HDMI molecule in individual nano-wells (Figure 1C, Figure S1, details in STAR Methods). This resulted in ∼80% of the nano-wells receiving a unique HDMI spatial barcode. The resulting data from the HDMI sequencing was processed, whereby HDMI sequences and their intra-tile coordinates were extracted from demultiplexed fastq files and stored in binary files for downstream usage (further details in STAR Methods). The preliminary HDMI sequencing quality, including the base composition was assessed prior to downstream steps (Figure S2)

After the HDMI cluster generation, the flow cells were subjected to a series of enzymatic treatments to prepare them for spatial transcriptomics profiling. Firstly, to expose the RNA capture site of the HDMI clusters, overnight incubation with DraI restriction enzyme was performed. Subsequently, the flow cell was treated with a cocktail of Exonuclease and Phosphatase to remove the leftover sequencing adapters in any nano-wells not containing a HDMI strand (Figure 1C).

Next, the flow cell was prepared for spatial transcriptomics capture (Figure 1D). The flow cell consists of three sandwiched layers: an upper thin glass layer, a gasket layer providing clearance for the fluidic flow and a lower thick glass layer (Figure 1D). The inner surfaces on both the thin and thick glass layers have functionalized nano-wells and both surfaces are used during sequencing. (Figure 1D). For cutting Nova-ST chips from the large NovaSeq S4 flow cells, we developed two different strategies. The first strategy is manual, where we begin by separating the thin and thick glass layers by gently prying between them with a fine edged scalpel blade. Once separated, the glass layers are scribed with a portable CNC diamond tipped scribing tool (Figure 1D, details in STAR Methods), along a rectangular grid of 1cm x 1cm (Figure 1D). After this, standard glass manipulation tools were used to break the large piece of glass along the scribed lines, separating it into individual Nova-ST spatial chips. A fully successful disassembly yields 36 thick and 36 thin, 1 cm x 1 cm Nova-ST chips from a single NovaSeq S4 flow cell (Figure 1D). The manual cutting strategy, particularly for cutting the thicker sections can be quite difficult and can result in uneven cuts.

To overcome this challenge, we also developed an automatic glass cutting strategy. In this method, the whole NovaSeq S4 flow cell is diced using a wafer dicing instrument (DISCO DAD 3220) into rectangular grid patterns of any desired dimensions (Figure 1E). The DAD 3220 employs a high-power, high-speed spindle (1.5 kW, 40,000 RPM) fitted with an extremely thin diamond blade that physically saws the glass substrate while producing cut-grooves in the range of just 200 µm. This dicing method provides a robust and very consistent workflow for producing Nova-ST chips compared to the manual cutting workflow. In the dicing method, the separation of the thin and thick glass layers is performed after dicing (Figure 1E) resulting in significantly reduced breakage and wastage. Following either cutting method, the Nova-ST chips were prepared for RNA capture after a series of washes and chemical treatment steps (details in STAR Methods, Figure 1E, Figure S1). Due to the flexibility of dicing, any chip dimension can be created, in this work we prepared several different chip sizes (5 mm x 8 mm; 6 mm x 8 mm and 10 mm x 8 mm) (Figure 1E).

### Implementation of spatial transcriptomics workflow

To test Nova-ST chips, we developed a spatial transcriptomics workflow for analyzing 10 µm fresh-frozen tissue sections. Here, we present spatial analysis results of several mouse brain coronal sections. The cryosectioned coronal regions were placed on the functional RNA capture area of the Nova-ST chips (Figure 2A). To perform tissue morphology registration for subsequent downstream alignment of the spatial data, hematoxylin and eosin (H&E) staining was performed on methanol fixed tissue. The spatial RNA footprint of the tissue was created by enzymatic permeabilization, RNA capture and first strand synthesis during reverse transcription on the HDMI functionalized surface of the Nova-ST chip (details in STAR Methods, Figure 2A, Figure S3). Subsequent second strand synthesis and denaturation followed by the PCR amplification yielded libraries ready for next generation sequencing (details in STAR Methods, Figure 2A & Figure S3). Paired end sequencing these libraries resulted in information about the spatial coordinates via the HDMI barcode and the gene expression corresponding to individual spatial barcodes (details in STAR Methods).

**Figure 2:**
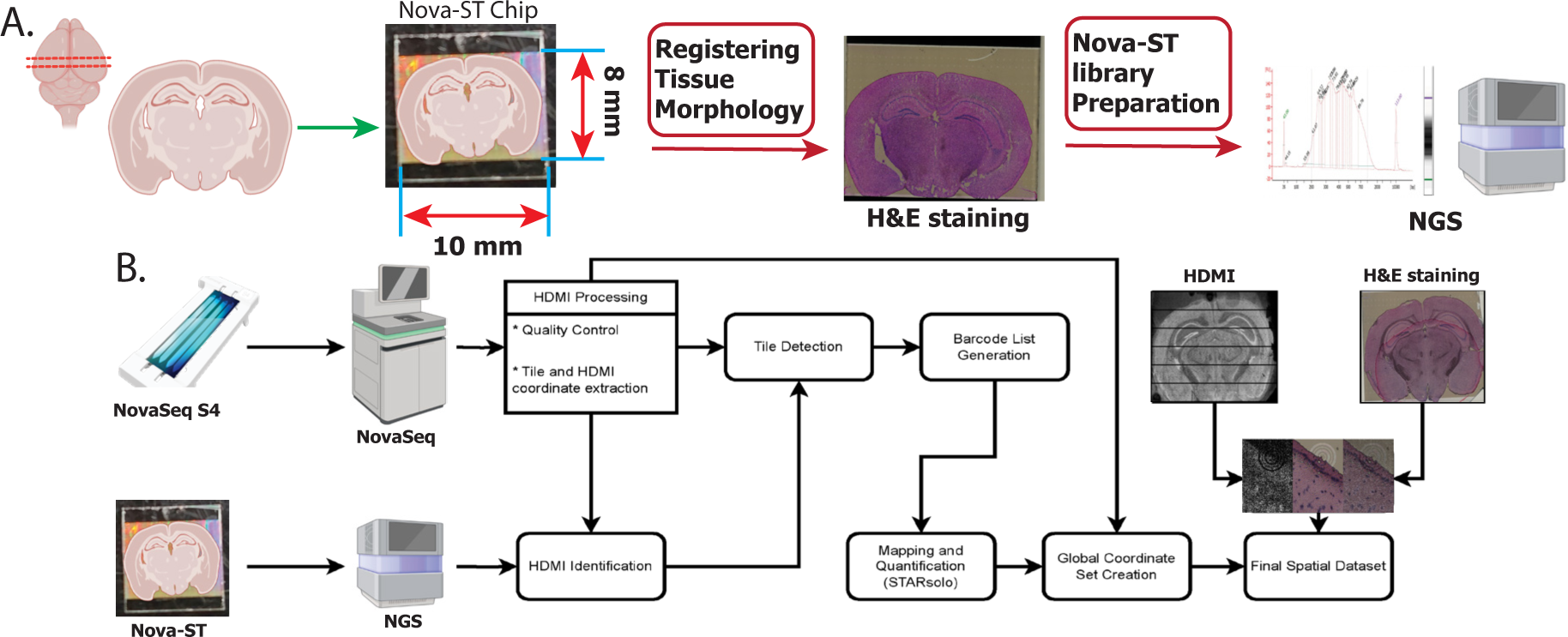
Overview of the spatial profiling with Nova-ST platform. A. Experimental steps in spatial transcriptomics profiling of a tissue with Nova-ST. Illustration of the mouse brain coronal sectioning and positioning on the Nova-ST chip. This tissue is then stained with H&E for the subsequent downstream tissue registration used during the spatial analysis. After the final library preparation, the quality of the library is analyzed and sequenced. B. A flow chart showing a schematic overview of the data generation and pre-processing steps in the Nova-ST pipeline.

During initial HDMI sequencing on the NovaSeq instrument, the sequencer images the flow cell in small contiguous sections called tiles, and the coordinates of each sequenced cluster is reported within the context of these tiles rather than the whole flow cell. As these tiles are not physical features on the flow cell, it is a priori not known which Nova-ST chip contains which tiles. Therefore, in the Nova-ST analysis pipeline developed here (Figure 2B) we begin with identification of these tiles by processing a small portion of the reads from the spatial transcriptomics library and comparing them with a subset of the barcodes sequenced per tile during HDMI generation. Next we extract all barcodes, as well as their spatial coordinates from the identified tiles to create a barcode whitelist that is then used for gene expression quantification using STARsolo^11^. (see STAR Methods, Figure 2B).

The NovaSeq flow cells also contain a pattern of physical fiducial markings that are visible under a microscope, as well as visible in the spatial coordinates due to their lack of nanowells, and therefore lack of HDMIs. Using the known distance between these fiducials (measured with light and electron microscopy), we updated the coordinates of HDMIs within each tile, placing them all within a global space of the Nova-ST chip. As this space is grounded in real measurements, we were able to align H&E images of the tissue section to the transcriptomic data using a simple affine transformation based solely on the fiducial markers (Figure 2B).

### Performance assessment of Nova-ST

The final HDMI footprint of the tissue indicated high spatial resolution (Figure 3A, 3B) also shown by the inset showing a zoomed-in area. We performed seven replicates across two batches of Nova-ST chips created from two separate NovaSeq S4 flow cells, replicates 1-5 are from one flow cell (FC1) and replicates 6 & 7 are from a second flow cell (FC2) (Figure 3C). The quality of the Nova-ST libraries is consistent between the chips derived from different flow cells and also between the thin and thick glass sections of the flow cells (Figure 3A-C). One random replicate was chosen for deep sequencing (∼1.2 billion reads), while multiple shallowly sequenced replicates clearly indicate the robust and consistent performance of the workflow (Figure 3A-C). The spatial resolution of the RNA footprint is similar between chip replicates from the same flow cell and between chips derived from different flow cells (Figure 3A, 3B). The performance, consistency and reproducibility between different Nova-ST chips derived from both the inter- and intra-S4-sequencing flow cells is also evident from the quality metrics of the spatial data (Figure 3C).

**Figure 3:**
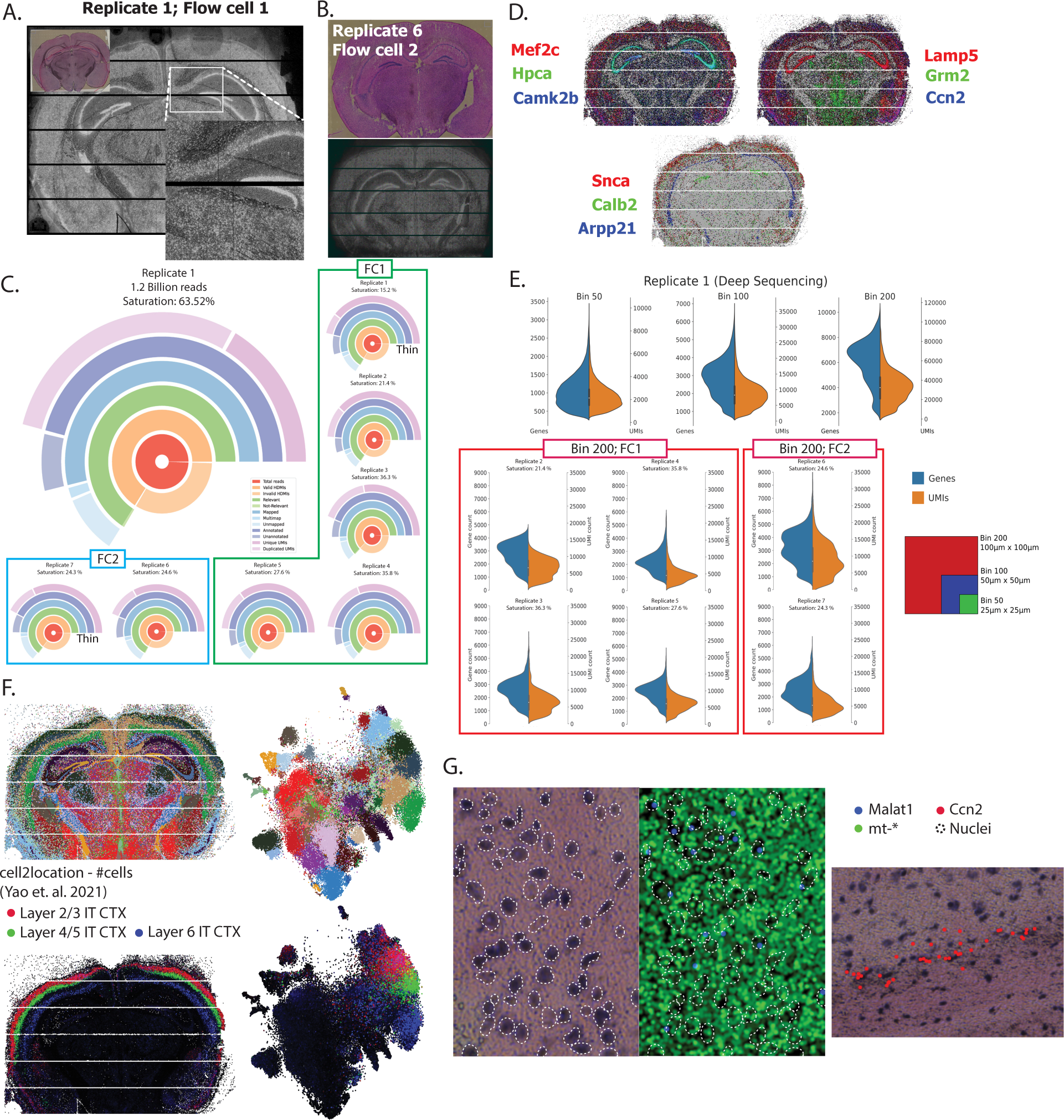
Performance of Nova-ST platform using mouse brain section. A. Spatial HDMI footprint of the transcripts captured from the mouse brain section in the deeply sequenced sample (DS) – replicate 1 (FS1). H&E stained tissue is present in the inset. B. Spatial HDMI footprint of the transcripts captured from the mouse brain and the H&E staining of the tissue of replicate 6 (FS2). C. Circle plots showing various metrics from the Nova-ST libraries, including mapping percentage, number of annotated reads and sequencing saturation, for all the mouse brain section replicates analyzed in this work. Most categories are self-explanatory, relevant, and non-relevant have no counterpart in Nova-ST data but are included for consistency with Stereo-seq samples. D. Spatial gene expression mapping of several region specific gene markers visualized with SCope^16^. E. Violin plots of both number of genes detected and number of UMIs detected at bin 50, 100 and 200 for the deeply sequenced replicate and bin 200 for all the remaining shallow sequenced replicates. F. t-SNE and spatial visualizations for Nova-ST DS showing Leiden clustering. Cell2location predictions for different layers in the cortex are also displayed. G. Two areas of tissues showing i) Malat1 expression localized to cell nuclei, distinct from mitochondrial reads and ii) Ccn2 expression limited to cortical layer 6b.

The data was binned into contiguous chunks of 25 µm x 25 µm (bin50), 50 µm x 50 µm (bin100) and 100 µm x 100 µm (bin200), and pre-processing was performed (see STAR Methods). Figure 3D shows the spatial expression of several marker genes, each localized in the correct brain region, such as Mef2c in the cortex; Hpca and Camk2b in the hippocampus; and Ccn2 in cortical layer 6b.

Next, we assessed the reproducibility of Nova-ST by comparing quality metrics across the seven replicates (six at lower sequencing depth [77-140 million reads]) (Figure 3B). The percentage of reads with valid barcodes that were also successfully mapped, ranged between 77.3% and 85.1% and the percentage of unique UMIs ranged between 36.48% and 78.60% (depending upon sequencing depth). For the three different bin sizes, bin50, bin100, and bin200, we detected a median of 994, 2821, and 6318 genes with non-zero counts; and 2131, 8268 and 32317 UMIs per bin for the deeply sequenced sample respectively; for the shallowly sequenced samples we detected a median of 263, 915, and 2594 genes, with a median of 393, 1613 and 6233 UMIs respectively (Figure 3E and Figure S4). Comparing total counts per gene across the filtered datasets at bin50 shows high reproducibility between samples (Figure S5, Table S2).

Unbiased clustering of bin50 data identified individual layers in the mouse cortex as well as subtypes within the hippocampus (Figure 3F). This resolution also allowed the identification of clusters of cells spanning just 1-2 cell layers, such as layer 6b neurons which formed a separate cluster (Figure S6) and agree with Ccn2 (a layer 6b marker gene), expression shown in Figure 3D and Figure S6. To test whether the unbiased clustering corresponded to known cell types, we used cell2location^12^ to map an independent scRNA-seq data set of the mouse cortex and hippocampus to the binned Nova-ST data^13^. This correctly identified the location of all annotated scRNA-seq cell types within the cortex and hippocampus, and cell types overall corresponded with the unbiased clustering (e.g., Layer 2/3, Layer 4/5, and Layer 6 intra-telencephalic (IT) neurons shown in Figure 3F). Next, we assessed RNA diffusion during tissue permeabilization. For this, we localized mitochondrial mRNAs and nuclear RNA (Malat1) and found these transcripts to be largely non-overlapping, suggesting low levels of RNA diffusion (Figure 3G). This was also the case with cortical layer 6b in the cortex where the expression of the Ccn2 gene is localized to the 1-2 cell layer thick cortical region (Figure 3G).

### Comparison of Nova-ST with existing technologies

Finally, we compared the performance (% of usable reads), the sensitivity (UMI and gene counts), and the specificity (correct localization of scRNA-seq cell types by cell2location^12^) of Nova-ST with Stereo-seq. To this end, we performed a Stereo-seq experiment on a comparable mouse brain section, and down-sampled the Stereo-seq data to the same depth as the deeply sequenced Nova-ST run. Nova-ST libraries have similar metrics to Stereo-seq libraries with regards to mapping % and annotated reads (83.51% and 73.04% [Nova-ST] vs 77.9% and 62.55% [Stereo-seq]) (Figure 3C, 4A). Nova-ST shows a lower percentage of recovered barcodes, a large portion of these are explained by sections of the chips that are not sequenced in the initial NovaSeq run but are still functional (visible as rows with no counts in Figure 3A). Nova-ST has superior complexity, allowing more genes/UMIs to be detected at lower sequencing depths and greater information obtained in deeper sequencing (25.54% [Nova-ST] versus 9.85% [Stereo-seq] unique UMIs at equal depth) (Figure 3C, 4A). At a bin size of 200, Nova-ST detects a median of 6318 genes compared to 4092 genes with the Stereo-seq platform at the same bin size (Figure 3E, 4B).

**Figure 4:**
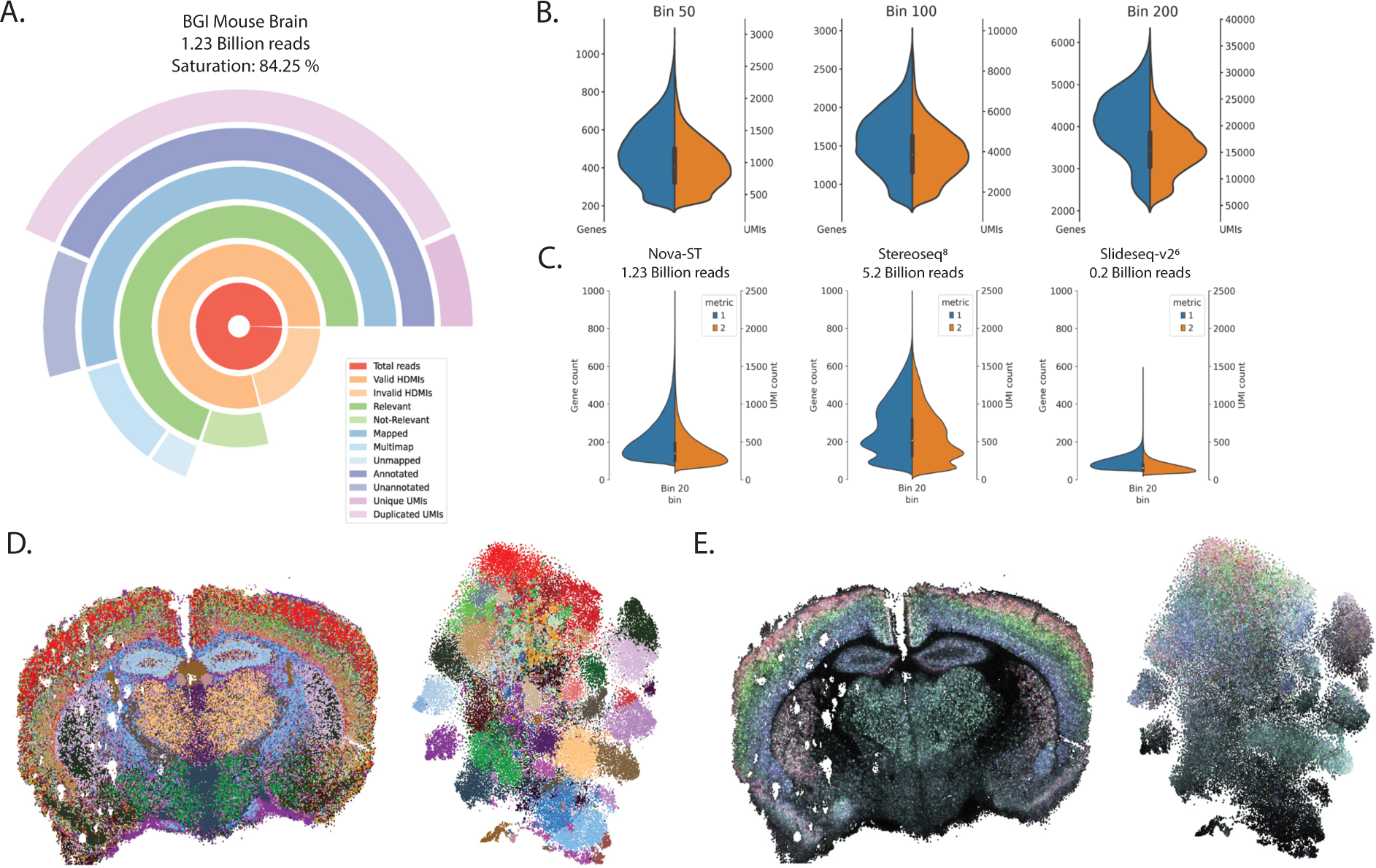
Comparison of Nova-ST with other sequencing based spatial transcriptomics technologies. A. Circle plot displaying the sequencing metrics of a Stereo-seq coronal mouse brain section, down sampled to 1.23 billion reads to match the Nova-ST DS sample. B. Violin plots of bin50, bin100 and bin200 datasets showing the distribution of the number of genes and UMIs detected across bins for the in-house BGI Stereo-seq sample. C. Violin plots of gene and UMI counts across bin20 (10µm x 10µm) data from three technologies, Nova-ST (this study), Stereo-seq^8^ and Slideseq-v2^6^. D. Spatial and tSNE visualizations of Leiden clustering (resolution 4.0) on our in-house Stereo-seq data. E. Spatial and tSNE visualizations of cell2location predicted number of cells for cortical layer neurons (legend in Figure 3E.)

Further comparing with other methods, we also binned our Nova-ST data in 10 µm x 10 µm bins (bin20) to compare with previously published Slideseq-v2 data ^6,14^ and another publicly available Stereo-seq sample^8^ (Table 1, Figure 4C). We found that at a similar depth we out-performed our in-house Stereo-seq sample, we detect greater numbers of genes and UMIs than Slide seq-v2 (albeit with greater sequencing depth), and despite having over 4 times the number of reads, and covering half the amount of tissue, when comparing with the public Stereo-seq sample, we detect only ∼20% fewer genes.

**Table 1.** Sequencing metrics from various sequencing based spatial transcriptomics technologies at bin20 (10 µm × 10 µm).

| Sample | Area covered | Reads | Median Genes | Median UMIs |
| --- | --- | --- | --- | --- |
| Nova-ST | Entire coronal section | 1.23 billion | 199 | 349 |
| In-house Stereo-seq | Entire coronal section | 1.23 billion | 91 | 155 |
| Slide-seq-v2<sup>6,14</sup> | Hippocampus only | ~0.2 billion | 153 | 211 |
| Stereo-seq<sup>8</sup> | Half coronal section | 5.2 billion | 266 | 519 |

Open-ST^15^, another spatial sequencing technique developed independently by another group also demonstrates superior sensitivity compared to Stereo-seq. For Nova-ST, we detect a median UMI count of 294 per 100 µm^2^ at a sequencing depth of ∼15 million reads per mm^2^ of total sequenced surface (DS sample), which compares to the sensitivity reported by the Open-ST platform^15^. Thus, Nova-ST obtains more genes and UMIs than Stereo-seq at similar bin sizes, allowing us to decrease bin sizes in Nova-ST to achieve increased resolution (Figure 3E, 4B).

Finally, unbiased clustering of our in-house Stereo-seq sample at bin50 shows some spatial localization of clusters, although without the ability to clearly delineate cortical layers, nor subtypes in the hippocampus (Figure 4D). When performing the same cell2location analysis as described above with the Nova-ST sample, cortical layers are visible, however they are both diffuse, and contain high level of background signal across the tissue section (Figure 4E) when compared to the predictions on our Nova-ST data (Figure 3F).

## Discussion

Spatial analysis is becoming a common place method in many fields of biomedical research, including cancer biology, neuroscience, and developmental biology^4^. Sequencing based spatial barcoding workflows offer flexibility, ease of use and the ability to perform unbiased spatial transcriptomics on any desired tissue and species. Existing sequencing-based technologies have several limitations with spatial resolution, area for spatial capture and most importantly the financial feasibility of profiling large scale tissue sections. To overcome some of these challenges, we have developed Nova-ST, a workflow that uses spatially barcoded nano-patterned Illumina sequencing flow cells to perform spatial transcriptomics.

To enable large throughput spatial screening, we first evaluated different Illumina sequencing flow cells and chose the NovaSeq S4 flow cell for developing Nova-ST. The Nova-ST chips are prepared from randomly barcoded NovaSeq flow cells obtained from 1^st^-level HDMI sequencing. We devised two different cutting strategies to make the barcoded NovaSeq flow cell amenable for spatial sequencing, one based on manual manipulation of the separated flow cell glass layers, while the second method utilizes an automated glass dicing technique to cut the flow cell to desired dimensions. We found both methods to be robust, producing many Nova-ST chips from a single flow cell, yet the automated method proved more sensitive allowing for further customized sizes, as well as minimal breakage & wastage. Furthermore, the automated dicing strategy also resulted in minimal damage or artifacts to the functional surface of the chips, enhancing the efficiency of the spatial capture and the quality of the data.

To demonstrate the performance of Nova-ST chips and benchmark against other spatial barcoding technologies, we developed a spatial transcriptomics workflow and used commonly used frozen coronal mouse brain sections. Nova-ST uses standard H&E staining for spatial registration of the tissue and for the subsequent downstream alignment of the spatial data. We developed a computational workflow that rapidly processes both the initial NovaSeq data and the spatial libraries to produce easy-to-use files for downstream processing and registration, while also providing multiple QC steps for assessing data and spatial library quality.

Spatial analysis on mouse brain sections indicated superior performance of Nova-ST, with high spatial resolution of transcript footprint of the tissue. The sequencing quality metrics indicated the Nova-ST transcriptome libraries to have a wide transcript diversity. The results also displayed consistency and reproducibility in spatial data between different chips derived from the same NovaSeq S4 flow cell as well for chips derived from different flow cells. Notably, the binned gene expression pattern of several gene markers was localized in the expected brain regions with near single cell resolution. Expression of the gene Ccn2, for example is localized to cells of the layer 6b of cortex and is just 1-2 cell layers thick; Nova-ST data recapitulates this across multiple independent replicates. Unbiased clustering of binned Nova-ST data robustly identified individual layers of the cortex and other brain regions, also further confirmed by the cell2location^12^ assignment using an independent scRNA mouse cortex dataset.

To compare and benchmark the performance of Nova-ST to other spatial technologies, we compared Nova-ST with in-house Stereo-seq data from the mouse cortex as well as published mouse cortex data from Stereo-seq^8^ and hippocampus data from Slide-seq v2^6,14^. When compared to the in-house generated Stereo-seq data, Nova-ST data indicated superiority in performance with higher gene detection sensitivity and higher ratios of the genes/UMIs detected at lower sequencing depths, facilitating greater sequencing economy. When compared to other publicly available spatial datasets derived from other spatial sequencing technologies Nova-ST displays higher or comparable performances.

Open-ST^15^ is another independently developed spatial transcriptomics technique based on the NovaSeq 6000 S4 flow cell and is very comparable to the Nova-ST workflow. Nevertheless, there are some key differences between the two implementations. Firstly, on the experimental side, the automatic dicing in Nova-ST provides robust chip cutting with fewer artifacts. Both the manual and automated cutting allow Nova-ST to use larger chip sizes without causing the glass to crack, compared to Open-ST (1 cm x 1 cm versus 4 mm x 3 mm). Secondly, on the computational side, the Nova-ST pipeline aligns the spatial data and H&E staining in a single step, whereas in Open-ST, each tile is aligned with the H&E image individually. This reduces the risk of producing alignments that are not consistent with real-world distances. In addition, Nova-ST generates a whitelist of barcodes for each tile based on the original HDMI sequencing, which is matched with barcodes discovered in the RNA sequencing (allowing for 1 mismatch). In Open-ST, all barcodes present within the RNA data are counted without comparison to a whitelist; this may result in decreased sensitivity when errors are introduced during sequencing. Overall, these are technical choices; and despite these differences, the Open-ST implementation confirms the QC and benchmarking statistics we observed with Nova-ST, suggesting that spatial transcriptomics on Illumina chips provides robust, cost-effective, and sensitive data.

Both these implementations still have limitations. A first limitation is that they require access to an Illumina NovaSeq 6000 instrument. For labs with no direct access to such a sequencer, it should be possible for a sequencing core facility to generate a HDMI barcoded flow cell and transport the chip in cold pack shipment. The flow cells are quite stable and can be stored at 4*C for at least 2 weeks. We recommend shipping the flow cell in its original shipping case to ensure safe transit. A second limitation is that chip cutting requires certain pieces of equipment, either a CNC equipped with a diamond-tipped scribing tool or a wafer dicing instrument for Nova-ST, or a 3D printed guide and diamond-tipped scribing tool for Open-ST. Of these, the manual scribing tool is most accessible, but may require some training on used Illumina chips, and particularly the thicker glass section is prone to breaking or may result in uneven cuts or loss of chips. Downstream handling with unevenly cut chips can also be very challenging and prohibits certain tissue sizes and/or shapes being used, and manual manipulation can lead to some damage on the functional area of the chips. Therefore, we recommend automatic dicing when specialized instrumentation is available. A third limitation is the 3’ tag sequencing. Like any other spatial barcoding ST assay that is based on 3’ tag sequencing, these techniques are not suitable for full length transcriptome analysis. However, we envision that with the slight modification of introducing a template switching oligonucleotide based first strand synthesis, full length transcripts can be obtained, which in combination with long read sequencing platforms such as Oxford Nanopore or PacBio can be used to interrogate splicing and isoform dynamics in a spatial context. A fourth limitation, particularly for Nova-ST, is that the data analysis is currently based on a binning strategy, combining contiguous regions of clusters together, this aggregation can lead to loss of single cell resolution. Nuclei based cell segmentation could be used to alleviate this problem, likely with a small decrease in sensitivity. A more elegant solution currently being investigated is the use of cell boundary staining to demarcate the cell boundaries. Another strategy that we are investigating is to reduce the tissue section thickness (∼5 µm). This would further decrease the transcript contamination arising from multiple layers of cells present in thicker sections.

Despite certain limitations and future opportunities, we showed in this work that Nova-ST is an easy to implement and most importantly a cost-effective way of performing sequencing based spatial transcriptomics at a higher or equivalent sensitivity compared to competing commercial methods. Unlike the recently introduced 10X Genomics Visium HD, Nova-ST is species agnostic and can also be used for studying non-model organisms. Nova-ST costs (including the reagents for first strand synthesis & library prep) approximately 7.5 € per mm^2^ of spatial area, excluding cost of sequencing. This is around 17% of the list price (∼45 € per mm^2^) of both the BGI Stereo-seq and 10X Genomics Visium HD. Together, this shows that Nova-ST not only outperforms the nearest competitive technology in terms of its performance in spatial profiling, but also will pave a way for cost-effective profiling of large-scale spatial transcriptomics. Due to the open-source flexibility of Nova-ST, further development for integrating other multiomics modalities such as open chromatin (ATAC) or spatial protein expression becomes a possibility, thus allowing broader use of Nova-ST and adoption within the scientific community for larger scale tissue transcriptomics.

## Acknowledgements

We thank UZ Leuven Genomics Core team for supporting with the sequencing workflow and gifting spent sequencing flow cells for downstream processing and flow cell cutting and processing practice. We thank Dr. Pieter Baatsen from the Electron Microscopy platform at VIB KU Leuven Center for Brain and Disease Research for his help with the surface scanning of the NovaSeq chip with Scanning Electron Microscope. We would also like to thank Mr. Donald Raddoux from Elektronische Circuits en Systemen (ECS), KU Leuven for their invaluable help and support with the DISCO DAD 3220 wafer dicing platform. Parts of Figure 2 and the graphical abstract were created with BioRender.com This work is funded by the following grants to S.A.: ERC Advanced Grant (101054387_Genome2Cells); Michael J. Fox Foundation for Parkinson’s Research (Michael J. Fox Foundation) (ASAP-000430).

## Author contributions

S.P., K.D. & S.A. conceived and developed the Nova-ST workflow; S.P. & K.D. performed the experiments and data analysis with help from R.V., N.P., A.V. & N.C.; S.P, K.D. & S.A. wrote the paper with inputs from all the other authors; S.P. & S.A. provided supervision and guidance for the project.

## Competing interest

The authors declare no competing interests.

## STAR Methods

### Resource availability

#### Lead contact

Further information and requests related to protocol details, resources and reagents should be directed to the lead contacts, Suresh Poovathingal and Stein Aerts

#### Materials availability

This study did not generate new unique reagents.

#### Data availability

Raw mouse brain data has been deposited to NCBI’s GEO archive: Nova-ST: GSE256318

Stereo-seq: GSE256319

Accessory data and information on the Nova-ST workflow can be found at: https://nova-st.aertslab.org/

#### Code availability

All code has been deposited on GitHub at: https://github.com/aertslab/Nova-ST

### Experimental model details

Mice were maintained in a specific pathogen-free facility under standard housing conditions (temperature 20-24°C and humidity 45-65%) with continuous access to food and water. Mice used in the study were 6-8 weeks old and were maintained on 14 hr light, 10 hr dark light cycle from 7 to 21 hr. In this study, whole brain from male C57BL/6J was used. Animals were anesthetized with isoflurane, and decapitated. Brains were dissected and immediately snap-frozen in isopentane for 10 min. No wild animals were used in this study. Sex is not relevant for this study as we report a spatial transcriptomics technique development as main finding, therefore sex was not considered in the study design. The findings in this study apply to only one sex (male mice). All animal experiments were conducted according to the KU Leuven ethical guidelines and approved by the KU Leuven Ethical Committee for Animal Experimentation (approved protocol no. ECD P007/2021).

### Method details

#### Mouse tissue preparation for cryo-sectioning

All animal experiments were performed according to the KU Leuven ethical guidelines and approved by the KU Leuven Ethical Committee for Animal Experimentation (approved protocol no. ECD P007/2021). Six to eight weeks old mice (C57BL/6J) used in the study were kept on a 14 h light, 10 h dark–light cycle from 7:00 to 21:00. Brains were dissected and immediately snap-frozen in isopentane for 10 min. Afterwards, brains were embedded in Tissue-Tek OCT cryo embedding compound. Coronal cryosections (10 µm) were performed at CT=14°C, OT=11°C. The used area of mouse cryosections is in the somatosensory areas adjacent to the posterior parietal association areas. For each brain tissue, 5-10 OCT scrolls of 70 um section thickness were collected into DNA lo-bind 2 ml eppendorf tubes. Ice cold PBS was used for washing the tissue to remove the OCT matrix. Total RNA was extracted from the washed tissue sections using the innuPREP mini RNA kit (Analytik Jena; Cat. No. AJ 845-KS-2040250). The RNA quality was assessed using RNA Nano kit (Agilent). Only tissue with RIN > 7 was used for the spatial analysis.

#### Stereo-seq optimization of tissue permeabilization

Tissue optimization was performed using the Stereo-seq Permeabilization kit (Cat. No. 111KP118) and Stereo-seq chip set P (Cat. No. 110CP118) according to the manufacturer’s protocol (Stereo-seq permeabilization set user manual, Ver A1). Briefly, 4 permeabilization chips were removed from the storage buffer and washed with nuclease free water and dried at 37°C. Next, 4 consecutive 10 µm tissue sections were prepared from the tissue cryo-block and placed on the permeabilization chip, the tissue layer was thawed to attach it to the surface of the chip. After drying the tissue on a 37°C hot plate, the chip was then dipped into 100% methanol at -20°C and incubated for 30 mins to fix the tissue. Post fixation, the tissue permeabilization test was performed on these chips by permeabilizing the tissue with PR enzyme prepared in 0.01N HCl (pH 2.0), at 4 different time points ranging from 6 mins to 30 mins. After the permeabilization, the chips were rinsed with 0.1X SSC buffer before reverse transcription. Reverse transcription was carried out at 42°C for 1 hour in dark. Tissue removal was performed at 55°C for 1 hour using the TR enzyme to clear the tissue before imaging. Fluorescence imaging was performed in the TRITC channel with 10X objective, following the imaging guidelines provided by the manufacturer (Guidebook for Image QC & microscope assessment and imaging, Ver A5). The optimal permeabilization time was assessed based on the strongest fluorescence signal with the lowest signal diffusion (crispness of the RNA footprint). Based on our assessment, we found the most optimal permeabilization time for the mouse brain to be 12 mins.

#### Stereo-seq spatial transcriptomics analysis

The spatial transcriptomics analysis was performed using the Stereo-seq Transcriptomics kit (Cat. No. 111ST114) according to the manufacturer’s protocol (Stereo-seq Transcriptomics set user manual, Ver A2). Briefly, as with the permeabilization analysis, the T-chip was removed from the storage buffer and washed with nuclease free water and dried at 37°C. Next, a 10 µm tissue section from a desired region of interest was prepared from the tissue cryo-block and placed on the T-chip and thawed the tissue layer to attach to the surface of the chip. After drying the tissue on 37°C hot plate, the chip was then dipped into 100% methanol at -20°C and incubated for 30 mins to fix the tissue. The fixed tissue was then stained using the Qbit ssDNA reagent (Thermo Cat. No. Q10212). Fluorescence imaging of the single stranded DNA staining was performed in the FITC channel with a 10X objective, following the imaging guidelines provided by the manufacturer (Guidebook for Image QC & microscope assessment and imaging, Ver A5). Prior to permeabilization, the ssDNA-stained image was also subjected to QC analysis using the imageQC software as per manufacturer’s recommendations. As with the permeabilization protocol, the tissue permeabilization was carried out with PR enzyme prepared in 0.01N HCl (pH 2.0) at 37°C. The optimal permeabilization time estimated from the tissue permeabilization analysis was used for the transcriptomics analysis. After washing the chip, reverse transcription mix was added to the chip and incubated at 42°C for at least 3 hrs. Tissue removal from the stereo seq chip was achieved by incubating the chip in the TR buffer at 55°c for 10 minutes. cDNA release and collection was performed by incubating the chip in cDNA release mix overnight at 55°C and the released cDNA was purified with Ampure XP beads (Beckman Coulter; Cat. No. A63882) using the manufacturer’s recommendation. After quality assessment using a bioanalyzer (Agilent), sequencing library preparation was performed using transposase assisted tagmentation reaction. Indexed PCR and library purification was performed to prepare the final sequencing library as per manufacturer’s recommendations. Final Stereo-seq libraries were sequenced on MGI/BGI sequencing platforms and were sequenced at the MGI Latvia sequencing facility.

#### HDMI sequencing

The HDMI generation in this work was done using the Illumina NovaSeq 6000 S4 kit; 35 cycles (PN: 20044417). In accordance with the original publication^10^, HDMI32DraI-32 ultramer (IDT technologies – standard desalted purification) was used for first level sequencing on the NovaSeq S4 flow cell to generate the HDMI array. The HDMI32DraI-32 ultramer (details of all oligonucleotide sequences used in this work is provided in Supplementary Table 1) was diluted to 1 uM concentration and the actual concentration of the oligonucleotide was titrated using qPCR to estimate the final concentration to be loaded for sequencing. Briefly, we used the Kapa Library Quantification kit (Roche, KK4824) to quantify the Oligonucleotides. Based on the concentration estimated from qPCR, libraries were denatured and loaded at a final concentration of 300 pM on the NovaSeq 6000 following the manufacturer’s instructions. Custom read primer Read1-DraI was also ordered from IDT technologies with PAGE purification^10^. The read primer was diluted to 0.3µM with HT1 buffer and loaded into the custom read primer 1 position in the NovaSeq reagent cartridge. The sequencing configuration used for reading the HDMI barcodes was 37(R1)-0(I1)-0(I2)-0(R2). At the end of the 34^th^ cycle, the instrument was manually aborted without initiating a post run wash. The S4 flow cell was then retrieved for immediate downstream postprocessing, it can also be stored safely at 4°C for at least 2 weeks. Users not having direct access to NovaSeq 6000 instrument can instruct sequencing facility to perform the HDMI sequencing step and transport the sequenced flow cell at 4°C. Prior to shipping, the inlet and outlet ports of the flow cell should be sealed using a PDMS biopsy plugs (see the details below) to ensure the liquids in the flow channels does not dry out.

#### Post sequencing processing of the flow cell

Sealing of the inlet and outlet for the flow channels was achieved by plugging them with 1.5-2mm Polydimethylsiloxane (PDMS) cylinders. To prepare these cylinders, the monomer and catalyst of SYLGARD™ 184 Silicone Elastomer Kit (Dow chemicals) was prepared in a 10:1 weight ratio. The components were mixed thoroughly and vacuum degassed. To polymerize, the mix was poured into a 3 cm petri dish and incubated at 80°C for 2 hours to complete the polymerization process. The PDMS slab was then extracted from the petri dish and wrapped into aluminum foil. 2 mm PDMS cylinders were prepared from this slab using a 2 mm biopsy punch (World Precision Instruments; Cat No. 504531).

The HDMI flow cells were then subjected to downstream enzymatic treatment. Firstly, the flow channels were cleaned with 200 µl of nuclease free water, NFW (Thermo Fisher; Cat No. 10977035). This was repeated for a total of three times. During each wash step, after filling the channels with reagents, a vacuum source (general vacuum pump, e.g. VWR, Cat. No. SART16694-1-60-06) was used to completely remove the reagents from the channel. Aspiration was continued until the channels became completely dry. To expose the RNA capture handle, the double stranded DNA was cut using restriction endonuclease DraI (NEB Inc. Cat. No. R0129L). The flow channels were first cleaned with 200 µl 1X rCutSmart™ Buffer. Then all flow channels were filled with 200 µl DraI reaction mix (1X rCutSmart™ Buffer, 2U/ul DraI Enzyme). Excess liquid overflowing from the outlet ports was aspirated using the vacuum pump. After making sure there were no air pockets trapped in the flow channels, the 2mm PDMS cylinder blocks were forced into the inlet and outlet ports to seal them using thin tipped forceps. The flow cell assembly was then placed into a humidification chamber (Nunc Square BioAssay Dishes; Thermo Fisher – Cat. No. 240835). For humidification, the flow cell was placed along with wet paper tissue and the petri dish was sealed using Parafilm. The flow cell was then incubated overnight at 37°C for the completion of the endonuclease reaction.

The flow channels were washed three time with 200 µl of NFW. After the vacuum aspiration of water from the flow channels, the channels were filled with 200 µl of 1X Exonuclease buffer (NEB Inc. M0293L). After the removal of the exonuclease buffer, the flow channels were filled with 200 µl of Exonuclease reaction mix (1X Exonuclease reaction buffer, 1U/µl of Exonuclease enzyme and 0.14 U/µl of Quick Calf Intestinal Phosphatase (NEB Inc. M0525L)). After aspirating the excess reaction mix from the inlet/outlet of the sequencing flow cells, the ports were sealed with fresh 2 mm PDMS cylinders. The reaction mix was then incubated at 37°C for 45 mins in the same humification chamber. After the completion of the exonuclease reaction, the HDMI flow cell assembly was retrieved, and the channels were washed three times with 200 µl of NFW. After each wash the liquid was completely withdrawn from the flow channels using a vacuum pump.

### Flow cell disassembly for Nova-ST chip preparation

#### Manual cutting strategy

The flow cell assembly was next placed into an oven at 50°C for 20 mins to dry the flow channels. The orientation of the top and bottom glass layers was identified with respect to the inlet and outlet ports of the sequencing flow cells. This is required for identifying the spatial location of the Nova-ST chips during the spatial transcriptomics analysis. The glass chip was then removed from the plastic housing by manually pulling out the plastic brackets that clamp down the inlet and outlet ports, releasing the glass chip. Then, using a fine scalpel, the gasket layer between the thin and thick glass surfaces (Fig. 1d) was scored gently to separate the thin and thick glass layer. This scoring must be done carefully without damaging the functional surface of the HDMI array. After gently prying and separating the glass layers, paper masking tape was glued to the back of the glass layers (3M, Cat No. 3M 201E 48MM), excess tape was trimmed off. In this work we have used the NOMAD 3 CNC milling machine from Carbide3D to score the glass surface into a 1cm x 1cm cutting grid (Fig. 1d). This CNC milling machine comes with a 130W spindle and has a working area of 200×200mm and 76mm in height. Less powerful machines can be used for the purpose described above. A diamond drag bit with a 90-degree tip, from the CNC milling machine manufacturer was used. The tip angle keeps the scoring as narrow as possible and ensures better penetration into the glass compared to a 120-degree tip. The bit is also equipped with a spring inside to adjust the force. In this case, it was adjusted so that little pressure is applied to the glass, while the tool length measurement probe of the machine can still detect the bit.

The scoring pattern was created using the machine suppliers dedicated software, Carbide Create. The glass plates dimensions were defined in the software as well as the desired scoring pattern. The scoring depths was then adapted according to the glass thickness and the direction. The thicker glass plate (1.2mm thick) was scored with a 0.6mm depth in the width direction (shortest side) and 0.2mm depth in the length direction. For the thin section (thickness 0.3mm), depths of 0.4mm and 0.1mm were used. It is important to know that the depth of cut defined in the software is not the actual depth. The actual depth differs due to the spring that retracts at the glass contact. This explains the higher depth of cut defined compared to the glass thickness. Each score was performed with a single pass of the tool.

Once the pattern was defined, the machine code (Gcode) was sent to the CNC machine via Carbide Motion, another software supplied by the manufacturer and was used to control machine movements. The plates were clamped to the table for scribing. After scoring the glass layers, the cutting was performed by using the glass running pliers (SPEEDWOX). To reduce the damage caused by the pliers on the functional surface, rubber tips were used. Before using the pliers, the pliers and the rubber tips were wiped with RNA & DNAZap followed by cleaning with 100% ethanol to ensure they were free of contamination. For cutting and preparing the Nova-ST chips, the pliers were used to cut the vertical score lines, by applying gentle pressure in the middle of the glass layers along the score line. After breaking the chips along the vertical score line, the masking tape was cut using a scalpel or razor. This was followed by breaking the glass chips along the horizontal score lines to produce the 1cm x 1cm Nova-ST chips. The Nova-ST chips were then pried out of the masking tape using forceps and the chips placed into 24 well plates. The location of the chip was recorded on the wells. This was repeated across the whole flow cell to produce 96 1cm x 1cm Nova-ST chips from both the thin and thick layers of the NovaSeq chip. In our experience breaking and preparation of the Nova-ST chips from the thick glass layer is non-trivial and sometimes getting perfect cut along the score lines is difficult. It’s highly recommended to practice the cutting on trial flow cells before attempting on the HDMI flow cells. The Nova-ST chips in the 24 well plates were then subjected to following steps to remove the second strand and to store them for long term. The Nova-ST chips were washed 3X times with 0.1N NaOH. For each wash the chips were incubated in the caustic solution for 5 mins, to ensure efficient denaturation of the second strand. Each of the Nova-ST chips were then washed 3X times with 1 ml nuclease free water followed by 2 times wash with 1 ml of 1X TE buffer (IDTE solution; IDT, Cat. No. 11-05-01-09). After the last wash, the chips were stored in IDT 1X TE buffer for long term storage at 4°C.

#### Automatic dicing strategy

After the final wash of the flow cell with NFW, the water in the flow channels was retained and the flow channels were sealed with PDMS plugs. The orientation of the top and bottom glass layers was identified with respect to the inlet and outlet ports of the sequencing flow cells, this is required for identifying the spatial location of the Nova-ST chips during the spatial transcriptomics analysis. The glass chip was then removed from the plastic housing by manually pulling out the plastic brackets that clamp down the inlet and outlet ports, releasing the glass chip. The NovaSeq flow cell was mounted on dicing tape which has a sticky backing that holds the flow cell on a thin sheet metal frame, to prepare for the dicing process. The thick glass side of the flow cell was glued on to the adhesive film. To reduce the adjustment for alignment in the dicing machine, the flow cell was glued on to the dicing tape, pre-aligned. The metal frame with flow cell was fixed to the dicing stage. The x-,y- and θ alignment was performed to align the flow cell. The origin for the start of the dicing was set. Cutting speed was set to 1 mm/s and dicing process was initiated. Firstly, the dicing was performed along the length of the NovaSeq flow cell and it was cut into 1 cm thick slabs. Without detaching the separated 1 cm slab from the dicing tape, the tape assembly was rotated by 90°, and the dicing was repeated along the width of the NovaSeq flow cell, cutting it into the desired dimensions. To ensure the glass layers of the flow cell were diced completely through, the score pattern on the dicing tape was checked. If there is no score pattern on the dicing tape, the NovaSeq chip has not been diced properly. After the dicing steps were completed, the diced NovaSeq flow cell was removed from the instrument and the dicing tape was cut out to retrieve the diced NovaSeq flow cell. With help of a sharp forceps Nova-ST chips were gently removed from the adhesive dicing tape. The Nova-ST chips at this stage still consist of two layers, the thin and thick sections still bonded by the middle gasket. A fresh razor was slid between the thin and thick sections and was used to gently pry, separating the chips. A gentle push is sufficient to separate the layers. Care should be taken to not disturb/damage the functional surface of the Nova-ST chips. Once the layers have been separated, the separated chips were placed on a paper towel with the functional surface facing up. After separating a batch of chips, using a sharp forceps, the chips were transferred to the respective 24 well plates, with the functional surface of the chips facing upwards. The Nova-ST chips in the 24 well plates were then subjected to following steps to remove the second strand and to prepare them for long term storage. The Nova-ST chips were washed 3X times with 0.1N NaOH. For each wash the chips were incubated in the caustic solution for 5 mins, to ensure efficient denaturation of the second strand. Each of the Nova-ST chips were then washed 3X times with 1 ml nuclease free water followed by 2 times wash with 1 ml of 1X TE buffer (IDTE solution; IDT, Cat. No. 11-05-01-09). After the last wash, the chips were stored in IDT 1X TE buffer for long term storage at 4°C.

#### RNA quality assessment

Prior to the tissue optimization and spatial transcriptome analysis on the Nova-ST chips, every tissue analyzed in this work was subject to RNA quality assessment. In brief, 5-10 serial sections of 50-70 um thickness were cryo-sectioned from a region farther away from the region of interest. These serial sectioned OCT scrolls were put into a 2 ml lo-bind tube (Eppendorf Cat. No. 0030108078) and stored at -80°C. The tissue scrolls were washed with 1 ml of ice-cold PBS at 4°C. Total RNA extraction from the spun-out tissue was performed using innuPREP mini-RNA kit (Analytik Jen; Cat. No. AJ 845-KS-2040250). Manufacturer recommendations were followed to extract total RNA. Elution was performed in 30 µl of NFW. The quality of the total RNA was assessed using Pico RNA kit (Agilent)

#### Optimization of tissue permeabilization for Nova-ST workflow

Optimal tissue permeabilization for mouse brain samples analyzed in this work was estimated using the 10X Visium Spatial Optimization kit (PN 1000192). Briefly, serial sections of the tissue were sectioned from the OCT embedded tissue matrix and placed on the capture spots of Visium Spatial Tissue Optimization slide (10X Genomics, PN: 3000394). To estimate the optimal permeabilization time, pepsin (Sigma Aldrich; Cat. No. P7000) at a concentration of 1 mg/ml in 0.1N HCl (Fisher Scientific Cat. No. AA35644K2) was used with different incubation times (5, 10, 15, 20, 25, 30, 35 mins). The rest of the protocol was followed as per the manufacturer’s recommendations (10X Genomics; Visium Spatial Gene Expression Reagent Kits – Tissue Optimization; CG000238 Rev E) to determine the most optimal time for tissue permeabilization. Imaging was performed using a Nikon NiE upright microscope equipped with a 10x Plan Apo lambda 0.45 air lens and a black and white sCMOS camera Prime BSI (Teledyne Photometrics). The setup was controlled by NIS-Elements (5.42.04, Nikon Instruments Europe B.V.). TRITC was excited with 550nm (CoolLED pE-800) and collected with a 577-630nm emission filter. A large tilescan was acquired to cover the entire tissue and chip area using 10% overlap and a focus surface. Like Stereo-seq, the optimal permeabilization time was assessed based on the strongest fluorescence signal with the lowest signal diffusion (crispness of the RNA footprint). Based on our assessment, we found the permeabilization time of 27 minutes optimal for mouse brain sections.

#### Nova-ST workflow: Tissue preparation, permeabilization & first strand synthesis

As with Stereo-seq experiments, the OCT embedded tissues were sectioned to a thickness of 10µm, placed on the capture area of the Nova-ST chip and melted. If the samples were not immediately processed for transcriptome capture, the Nova-ST chip was re-frozen on cryoblock and stored in -80°C and in our experience the quality of the tissue is not impacted with 2-3 weeks of storage of the tissue section at -80°C.

Standard Hematoxylin and Eosin (H&E) staining procedure was used to stain the tissue. Briefly, the Nova-ST chip with frozen tissue section was taken from -80°C storage and immediately melted on a 37°C block for 1 min. The tissue was then fixed in methanol at - 20°C for 30 mins. Post fixation, the tissue was dehydrated by adding 150µl of isopropyl alcohol (IPA) and incubating for 1 min. After removal of the IPA, the Nova-ST chip was air dried for 3 mins (or until the chip is completely dried). 200 µl Mayer’s haematoxylin (Agilent, Cat. No. S3309) was added to the chip and incubated for 7 mins. Using a forceps the chip was washed in excess NFW. 150µl of bluing buffer (Agilent, Cat. No. CS702) was added to Nova-ST chip and incubated for 2 mins. The chip was again washed with excess NFW. The tissue was then treated with 200 µl of Eosin-Y buffered solution (10% v/v of Eosin-Y (Sigma, Cat. No. HT110216) in 0.45 M Tris Acetic acid solution pH 6.0) and incubated for 1 min. The chip was dried at 37°C for 5 mins (or until the water was completely evaporated) prior to imaging. Brightfield imaging was performed using a Nikon NiE upright microscope equipped with a 10x Plan Apo lambda 0.45 air lens and a color camera DFK 33UX264 (The Imaging Source, LLC). The setup was controlled by NIS-Elements (5.42.04, Nikon Instruments Europe B.V.). A large tile scan was acquired to cover the entire tissue and chip area using 10% overlap and a focus surface. Post imaging, the sample was immediately processed for the spatial transcriptomics workflow.

Pepsin reagent prepared in 0.01N HCl (pH 2.0) (1 mg/ml) was preheated in a 37°C oven. After H&E staining and imaging, using forceps, the Nova-ST chip was placed into a 3 cm petri dish. 300µl of prewarmed pepsin was added to the H&E stained tissue and permeabilization was performed at 37°C for the optimal permeabilization time estimated in the previous step. After the permeabilization step, the pepsin solution was blotted off from the Nova-ST chip and the chip was transferred into a 24 well plate. The permeabilization reaction was stopped by sequentially washing the chip with 0.1X SSC (20X SSC; Thermo Fisher; Cat. No: 15557044), followed by 300 µl of 1X RT wash buffer (1X Maxima h-Reverse Transcriptase buffer; Cat. No EP0753, 1U/µl Lucingen NxGen RNase Inhibitor; Cat. No. 30281-2). Finally, 300 µl of First Strand mix was added to the well (1X Maxima h-Reverse Transcriptase buffer, 1U/µl Lucingen NxGen RNase Inhibitor, 4% Ficoll PM-400; Sigma Aldrich Cat. No. F4375-10G, 1 mM dNTP; Thermo Fisher Cat. No. R1121, 10U/µl Maxima RTase). The wells were covered with multiple layers of square patches of Parafilm. The 24 well plate was then sealed and put into oven at 42°C for overnight incubation for first strand synthesis.

#### Nova-ST workflow: Exonuclease treatment

In this step, exonuclease treatment was performed on the Nova-ST chip to remove single stranded HDMI capture tags without the first strand product to avoid the undesired secondary downstream reactions. After the first strand reaction, the FSS mix was removed from and the Nova-ST chip and was washed with 300 µl of 0.1X SSC. The chip was washed with 300 µl of 1X Exonuclease I buffer before adding 300 µl Exonuclease reaction mix (1X Exonuclease reaction buffer, 1U/µl of Exonuclease enzyme). The reaction was incubated at 37°C for 45 mins.

#### Nova-ST workflow: Tissue clearance

After incubation, the tissue on the surface of the Nova-ST chips was cleared. In this step, the exonuclease reaction mix was removed from the well and 300 µl of Tissue clearance reagent was added to the well containing the Nova-ST chip (100 mM of Tris pH 8.0; Thermo Fisher Cat. No AM9856, 200 mM of NaCl; Thermo Fisher Cat. No. AM9760G, 2% SDS; Thermo Fisher Cat. No. 24730020, 5 mM EDTA; Thermo Fisher Cat. No. 15575020, 16U/µl Proteinase K; NEB Inc Cat. No. P8107S). The reaction was incubated at 37°C for 45 mins to complete the tissue removal reaction.

#### Nova-ST workflow: Second strand synthesis

Before the subsequent processing, clearance of the tissue was ensured from the surface of the Nova-ST chip. Then the Nova-ST chip was washed three times with 3 ml of NFW. The chip was them washed three times with 0.1N NaOH. During each wash, the chip was incubated in 0.1N NaOH for 5 mins to remove the mRNA strand. The Nova-ST chip was then washed three times with 0.1M Tris-HCl (pH 7.5) (Thermo Fisher; Cat. No. 15567027) followed by 3X times wash with NFW. Using forceps, the chip was transferred to another well in the 24 well plate. Before transferring excess water was blotted off from the bottom of the chip surface. The chip was then subjected to second strand synthesis reaction. 300µl of Second strand synthesis reaction mix (1X NEB Buffer 2, 10 mM RPE randomer, 1 mM dNTP, 0.5 U/µl Klenow Fragment (3’→5’ exo-); NEB Inc Cat. No. M0212L) was added to the well. The chip was incubated in the second strand reaction mix for 2 hours at 37°C.

#### Nova-ST workflow: Second strand product extraction, cleanup and random primer extension PCR

The Nova-ST flow cell was washed three times with 3 ml of NFW. Using forceps, the chip was transferred from the well to a 3 cm petri dish. Before transferring excess water was blotted off from the bottom of the chip surface. 90µl of 0.1 M NaOH was added to the surface of the Nova-ST chip, it was then incubated on the chip for 5 mins. After the incubation, the liquid was harvested into a DNA lo-bind 1.5 ml eppendorf tube. This process was repeated two additional times. After the final collection, the volume of the RPE collect was estimated and 0.28 times the volume of Tris 7.0 (Thermo Fisher Cat. No. AM9851) was added to the RPE collection to neutralize the reaction. After two minutes of incubation, the RPE product was purified using 1.8X Ampure XP beads as per manufacturer’s recommendation. The magnetic bead elution was performed with 44 µl of EB buffer (Qiagen Cat. No. 19086). 42 µl of elute was taken for the RPE PCR. PCR was performed on purified RPE product by adding the PCR mix (1X KAPA HiFi master mix; Roche Cat. No. KK2602, 1 µM RPE forward primer, 1 µM RPE reverse primer). The following RPE PCR program was used for the product amplification: 95°C – 3 mins, 14 cycles of {95°C – 30s; 60°C – 1 min; 72°C – 1 min}, final extension of 72°C – 5mins. The PCR product was then purified with 0.8X Ampure XP beads with elution in 40 µl of EB buffer.

#### Nova-ST workflow: Index PCR and sequencing

The RPE amplified library was quantified using the Qubit dsDNA Quantification kit (Thermo Fisher Cat. No. Q32851) and the size distribution of the RPE amplified library was estimated using High Sensitivity DNA kit (Agilent). Based on the measurements 10 nM RPE library dilution was performed using NFW. Final indexing PCR was performed by adding the PCR mix (1X KAPA HiFi master mix, 1 µM WTA forward primer, 1 µM WTA reverse primer and 2nM of RPE library). The following WTA PCR program was used for the product amplification: 95°C – 3 mins, 14 cycles of {95°C – 30s; 60°C – 30 s; 72°C – 30 s}, final extension of 72°C – 5mins. Two rounds of purification were performed on the amplified PCR product with 0.8X Ampure XP beads and the final elution in 60 µl of EB buffer. Sequencing of the Final Nova-ST libraries was performed on NextSeq 2000 sequencing platform. The concentration of the libraries were normalized to 2 nM using RSB buffer with Tween-20 (Illumina; Cat. No. 20512944). The 2 nM library was further diluted to 800 pM before loading to the instrument. The libraries were sequenced with the following sequencing specification: R1 = 34 bps; I1 = 8 bps; I2 = 8bps; R2 = 91 bps.

#### Data Analysis – HDMIs

Raw sequencing data was loaded into Illumina’s Sequence Analysis Viewer to first, broadly check that the base composition matches the expected sequence (Supplementary Fig. 2) and secondly, to identify any tiles which were not sequenced – these were noted for exclusion in later steps. Raw data was next demultiplexed using Illumina’s bcl2fastq, creating one set of fastq files per tile of the flow cell using the following command: bcl2fastq -R ${RUN_FOLDER} -o Demultiplexed/${TILE_NO} -r 1 -p 1 -w 1 --tiles s_<${TILE_NO}> --use-bases-mask=y32n* --minimum-trimmed-read-length=32 --write-fastq-reverse-complement. For each read in each fastq file, the barcode sequence was first checked against the expected degenerate sequence and then the tile number, X coordinate, Y coordinate and read sequence, were recorded and saved to disk. A small subset (10,000) of valid reads was saved to a separate file for later use.

As the X and Y coordinates obtained during the previous step were local coordinates and specific to the tile that each read came from, to properly reconstruct the spatial location of every read from the chip, these coordinates were put within a global context, we use the fiducial markers (concentric circles) present within each tile to align them to each other. This step can either be performed on a per-chip basis, or across all tiles from the HDMI generation, but was performed per chip for the data in this study. To do this, for each tile processed, a numpy array (max_x, max_y) was created and the value at each coordinate where a valid barcode is found was set to 1. These matrices were trimmed and reshaped to bin the data in 25×25 bins, simplifying processing. Matrices were normalized to a maximum of 255 and then converted to greyscale images using OpenCV. These images were inverted, denoised (cv.fastNlMeansDenoising, h=100) and thresholded (min 128, max 255) to extract and image of just the fiducial markers. Next, Hough circle detection was used at 2 different radii (1. minRadius=40,maxRadius=80, 2. minRadius=15,maxRadius=30) to detect the coordinates of the inner and outer fiducial circles within each tile and the centroid of each set was calculated when all (8) circles were identified successfully. Where all 8 fiducials were not identified, centroid coordinates were interpolated using the adjacent tiles. Distances between fiducial circles were measured in the H&E and electron microscopy images (in both rows/swaths and columns) and these distances were used to calibrate the scale of the coordinates extracted from the fastq files to nm. HDMI coordinates were then corrected per tile to place the fiducial circles the correct distance from each other in both directions, beginning with the top left tile of a Nova-ST chip. Swaths 1, 3 and 5 begin at the same position, swaths 2, 4 and 6 are offset by a single tile, and this was accounted for in this correction.

#### Data Pre-processing – Spatial Libraries

Barcodes from the first 1 million reads from read 1 of the RNA sequencing were extracted and compared with the subset taken in HDMI Basic Processing to identify the tiles in the Nova-ST chip used which have reads (i.e. were under the tissue section). All barcodes from the HDMI data of the tiles covered by the section were then extracted into a whitelist for STARsolo (2.7.10b), these were trimmed to 31bp to allow STARsolo to perform a Hamming error-correction. For each read of the spatial library, the UMIs present in read 2 (the first 9 base pairs) were extracted and appended to the corresponding read from read 1 and STARsolo was run with the following parameters: --soloType CB_UMI_Simple --soloCBwhitelist ${BARCODE_WHITELIST_FILE} --soloCBstart 1 --soloCBlen 31 -- soloUMIstart 32 --soloUMIlen 8 --soloBarcodeMate 0 --soloBarcodeReadLength 0 -- soloFeatures Gene GeneFull --soloCBmatchWLtype 1MM --soloUMIdedup 1MM_All -- soloCellFilter None --outSAMtype BAM SortedByCoordinate --outSAMattributes NH HI AS nM CR CY UR UY CB UB sS --readFilesIn ${READ_2} ${READ_1_PLUS_UMI} Matrices from STARsolo and the corrected coordinates were combined into a GEM file, a format used by BGI’s Stereo-seq pipeline, to enable the loading and binning functions of Stereopy, as well as consistent analysis of the two data types.

#### Data Analysis – Spatial Libraries

Stereo-seq matrices were loaded into Python using Stereopy (v0.6) at three different bin sizes, 50, 100 and 200, for Nova-ST, the following sizes were used 728, 1456, 2912, where each size corresponds to the same dimensions as the Stereo-seq bins. Following data loading, samples were further analyzed using Scanpy, in brief: Standard quality control metrics were calculated and bins with too few genes detected were removed (Bin 50/728: 75 Genes, Bin 100/1456: 250 Genes, Bin 200/2912: 500 Genes), bins were not filtered by mitochondrial percentage. Bins were normalized to a total of 10,000 counts and log transformed before highly variable genes were identified. A regression for total counts and mitochondrial percentage was applied and counts were scaled to unit variance with a mean of 0, counts above 10 were clipped to 10. A principal component analysis was performed, followed by neighborhood detection, UMAP generation and leiden clustering at several resolutions. H&E images were aligned to spatial data using the BigWarp Fiji plugin, selecting multiple fiducial circles visible in both the H&E and spatial data as landmarks and using an affine transformation.

For cell2location, the single cell mouse cortex and hippocampus data from Yao et al 2021 was loaded from the provided h5 files into an Anndata object and associated metadata was added. This object was then subsampled without replacement to contain half of the original cells to simplify training the cell2location model. A cell2location RegressionModel instance was created using the single cell data with external_donor_name_label as the batch key and subclass_label at the labels key, and the model was trained with 500 max epochs. The cell abundance estimations were exported using the following parameters: num_samples=1000, batch_size=2500. Next, both the spatial and single cell data were subset to only include genes present in both datasets and a Cell2location model was set up using the spatial data (at bin50), the exported reference cell states and N_cells_per_location=2. This model was then trained using the following parameters: max_epochs=30000, batch_size=None, train_size=1) and the final estimated cell abundances was exported with num_samples=1000 and batch_size equal to 1/10^th^ of the dataset size. Anndata objects were converted into loom files, including the spatial coordinates and cell2location prediction scores for visualization in Scope.

## Supplementary information

- Supplementary_Figures.pdf: Figure S1-S6
- Supplementary_Table1.xlsx: primer sequences
- Supplementary_Table2.xlsx: read counts

## Notes

https://www.ncbi.nlm.nih.gov/geo/query/acc.cgi?acc=GSE256318

https://www.ncbi.nlm.nih.gov/geo/query/acc.cgi?acc=GSE256319

### Competing Interest Statement

The authors have declared no competing interest.

### Summary of Updates

This version includes a new automated method for cutting the chips using a wafer dicing instrument, as well as new replicates generated with these chips. Several comparisons with more publicly available data were also added.

