## Supplementary Figures for "Nova-ST: Nano-Patterned Ultra-Dense platform for spatial transcriptomics"

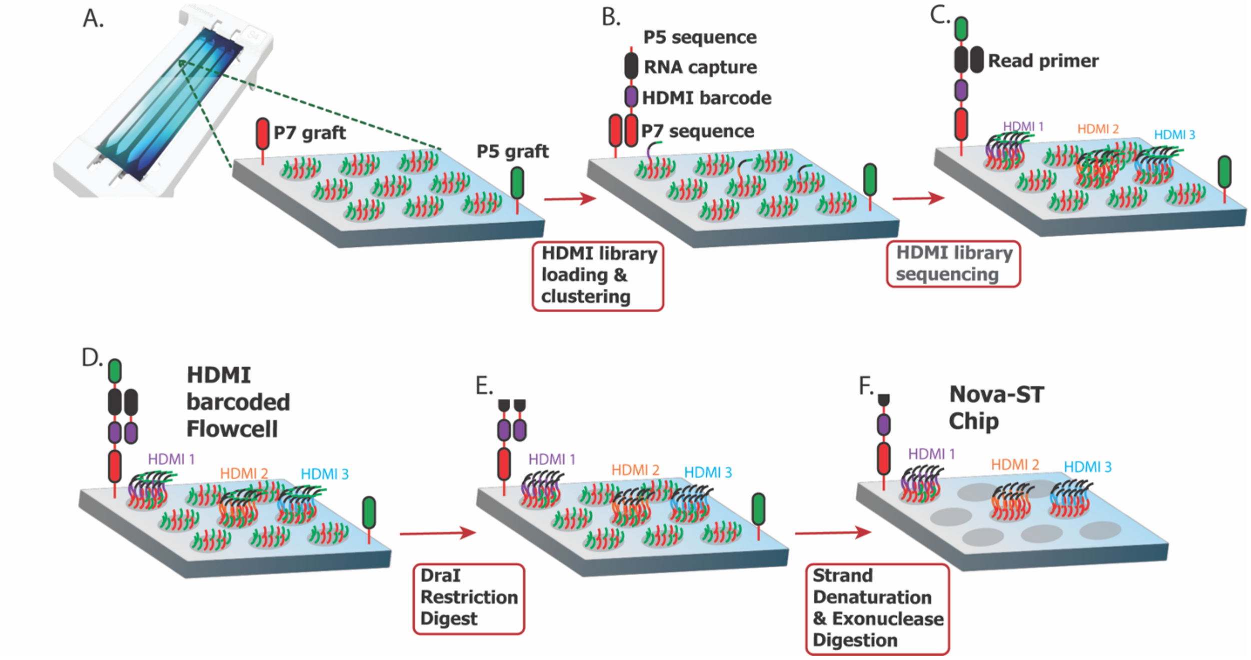


**Figure S1: Nova-ST chip preparation workflow.**

A.) Cartoon describing the nano well with the functionalized sequencing handles in the nano well of a NovaSeq 6000 S4 flow cell.

B-D.) HDMI sequencing workflow where the HDMI oligonucleotide (HDMI-DraI 32) undergoes annealing with the sequencing handles in the nano well. Then it undergoes isothermal bridge amplification in which several 1000 copies of single strand of the HDMI strand is created. During the sequencing, a custom read primer (Read1-DraI primer), reads the HDMI spatial barcode in the backbone of the HDMI oligonucleotide sequence.

E.) Post sequencing, the Dra-I restriction enzyme cuts the double strand sequence at the TTTAAA locus to free the capture domain.

F.) The double stranded DNA is denatured using caustic treatment to expose the RNA capture domain for spatial transcriptomics.

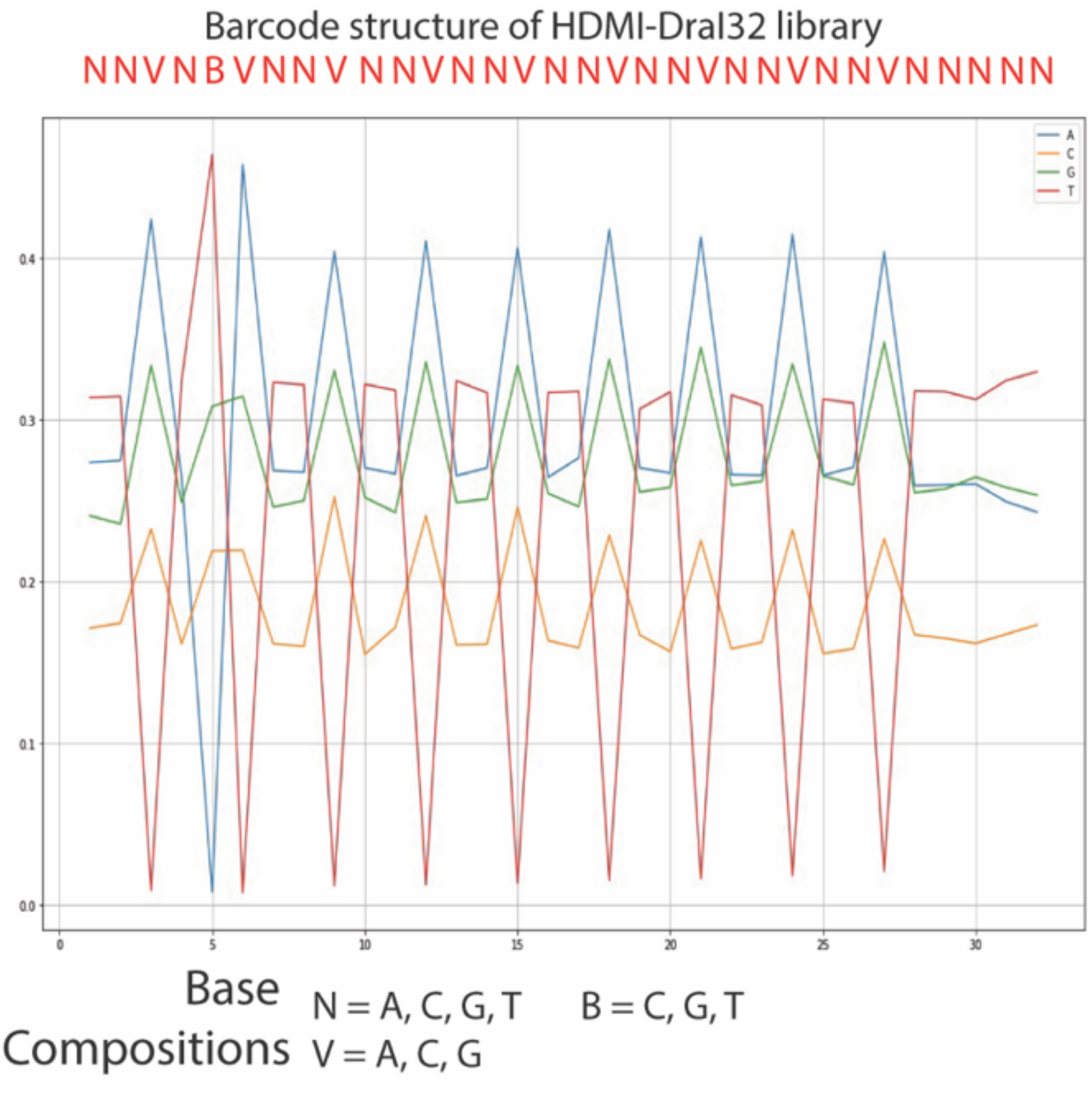


**Figure S2:** An example of base composition of the sequenced HDMI library compared to the actual sequence of the HDMI spatial barcodes in the HDMI-DraI32 ultramer.

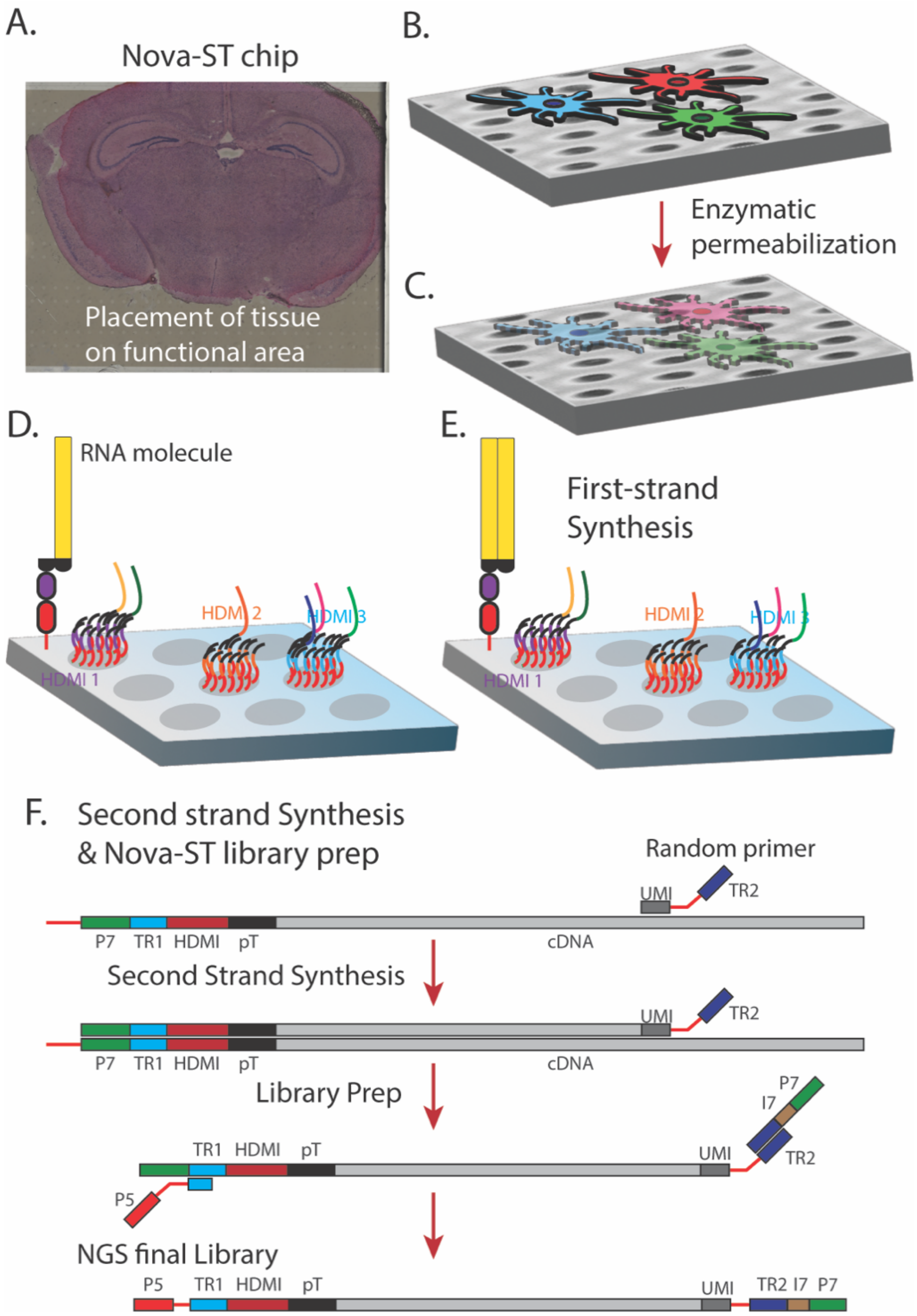


**Figure S3: Nova-ST spatial transcriptomics workflow.**
A.) Tissue overlaying on the functional surface of the Nova-ST chip.
B, C) Diagrammatic description of the enzymatic digestion of tissue on the surface of the Nova-ST flow cell.
D, E) mRNA capture from the permeabilized tissue followed by the first strand synthesis on the surface of the Nova-ST chip.
F.) downstream processing of the spatial library after the removal of the tissue from surface of Nova-ST chip, where the second strand synthesis is achieved by a random primer extension (RPE). The extended second strand product is denatured and the rest of the RPE product amplification and the indexed NGS library preparation is done in a tube.

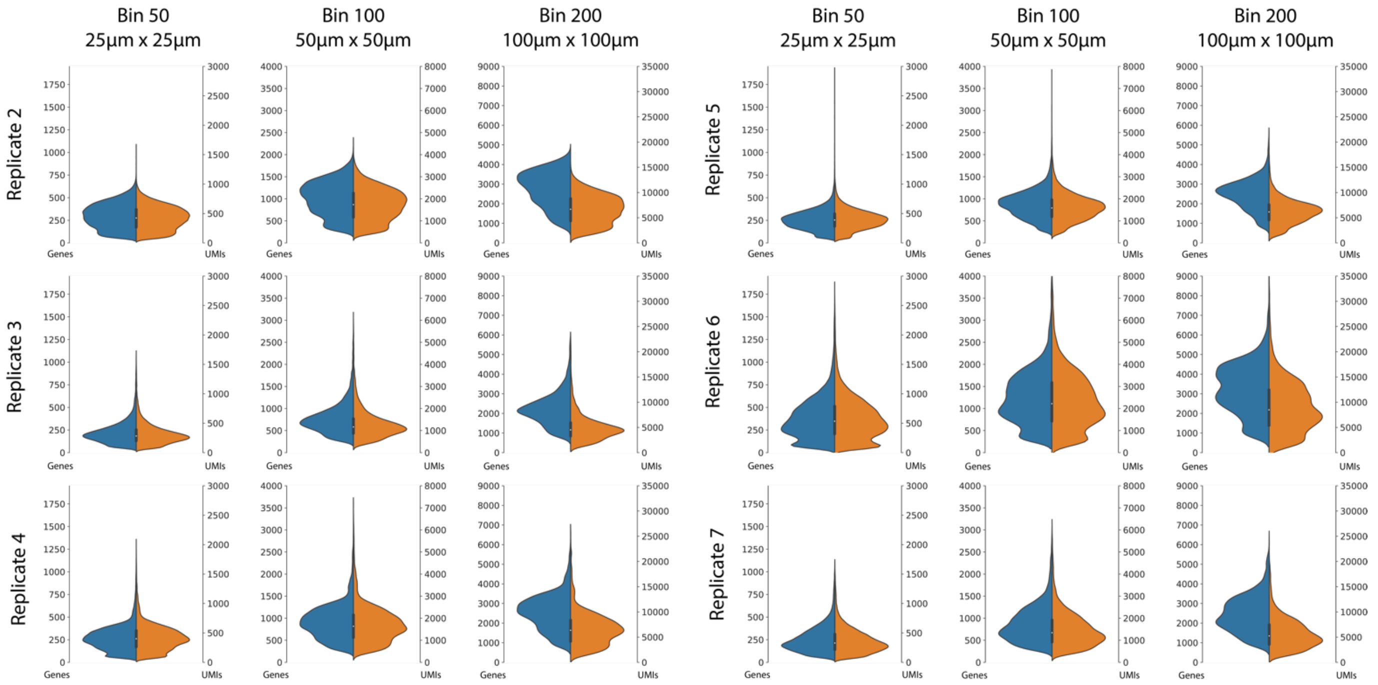


**Figure S4:** Gene (blue) and UMI (orange) distributions for all shallowly sequenced replicates at all three bin sizes (50, 100 and 200).

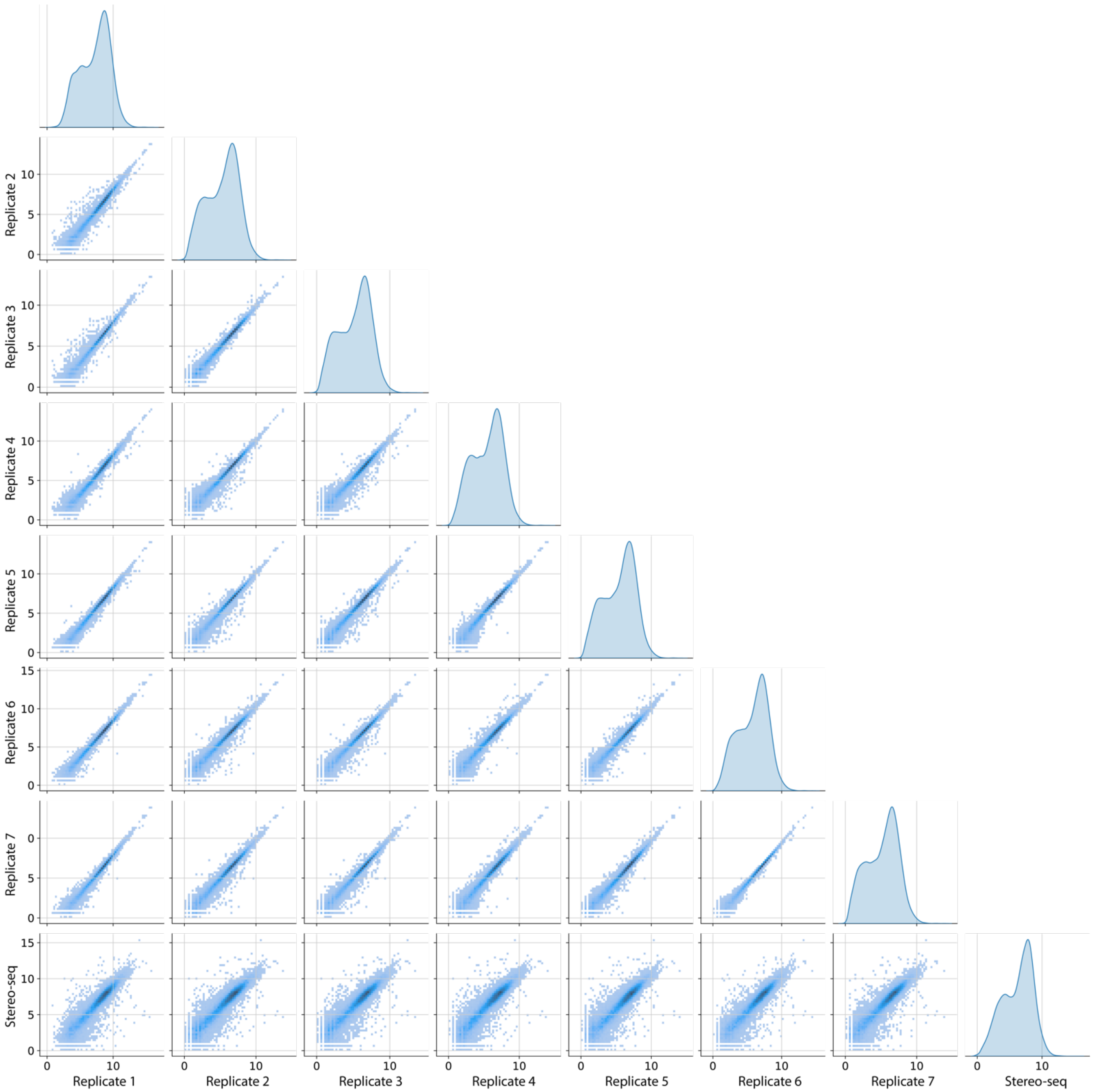


**Figure S5:** Pairwise comparisons of summed gene counts within each sample, log-log axes. Darker areas indicate higher genes density. Self-comparisons are excluded, only genes found in all samples are included.

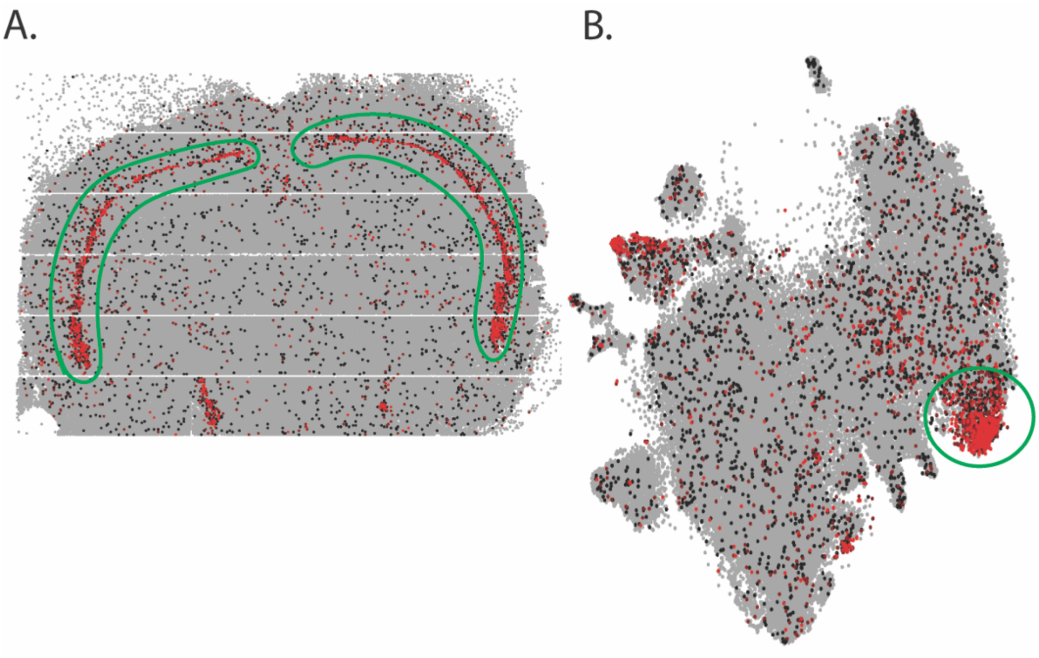


**Figure S6:** spatial visualizations and t-SNE (Leiden clustering) for Nova-ST DS data. Expression of Ccn2 localized to the cortex layer 6b and corresponding location of these bins in t-SNE map is highlighted by the green contours.
